# Draft genome of the Eutardigrade Milnesium tardigradum sheds light on ecdysozoan evolution

**DOI:** 10.1101/122309

**Authors:** Felix Bemm, Laura Burleigh, Frank Förster, Roland Schmucki, Martin Ebeling, Christian J. Janzen, Thomas Dandekar, Ralph O. Schill, Ulrich Certa, Jörg Schultz

## Abstract

Tardigrades are among the most stress tolerant animals and survived even unassisted exposure to space in low earth orbit. Still, the adaptations leading to these unusual physiological features remain unclear. Even the phylogenetic position of this phylum within the Ecdysozoa is unclear. Complete genome sequences might help to address these questions as genomic adaptations can be revealed and phylogenetic reconstructions can be based on new markers. Here, we present a first draft genome of a species from the family Milnesiidae, namely *Milnesium tardigradum*. We consistently place *M. tardigradum* and the two previously sequenced Hypsibiidae species, *Hypsibius dujardini* and *Ramazzottius varieornatus*, as sister group of the nematodes with the arthropods as outgroup. Based on this placement, we identify a massive gene loss thus far attributed to the nematodes which predates their split from the tardigrades. We provide a comprehensive catalog of protein domain expansions linked to stress response and show that previously identified tardigrade-unique proteins are erratically distributed across the genome of *M. tardigradum*. We further suggest alternative pathways to cope with high stress levels that are yet unexplored in tardigrades and further promote the phylum Tardigrada as a rich source of stress protection genes and mechanisms.

## Introduction

There is no life without water. Antony van Leeuwenhoek must have been well aware of this fact when in 1702 he collected some dry dust from a roof gutter. He was in for a surprise when he viewed the sample with one of his self-built microscopes. Soon after mixing with some clean water, he found tiny animals, which he called ‘animalcules’ [1]. Thus, seemingly dead animals came fully alive again after rehydration. In 1959, D. Keilin coined the term ‘cryptobiosis’, which can be triggered by low oxygen (anoxybiosis), low temperature (cryobiosis), high salt concentrations (osmobiosis) or desiccation (anhydrobiosis) [2]. One group of animals able to undergo cryptobiosis are Tardigrades (from latin tardus = slow and gradi = walk) [3]. These animals are about 0.1 to 1.2 mm in size with a peculiar shape reminiscent of bears. Accordingly, they have also been called ‘kleine Wasserbärchen’ (little water bear) in German. They were first identified at the end of the 18th century [4]. Today, more than 1,000 species are known [5,6]. As their German name already suggests, tardigrades are an aquatic life form and can only live covered by a water film. Still, most species inhabit terrestrial habitats like mosses and lichens which regularly fall completely dry. At these times, adults, juveniles and embryos [7] can only survive until the next rain period by changing from the active state into the anhydrobiotic tun state. As metabolic conversion of nutrients requires water, tardigrades in the tun state suspend life and do not age [8]. In this form, they survive being frozen [9,10], heated [11] and exposed to enormous levels of UV [12] or ionizing [13] radiation.

The first studies addressing the unique physiological peculiarities of tardigrades established different hypotheses regarding their underlying genomic basis. A genome wide analysis of the gene coding complement of the tardigrade *Hypsibius dujardini* found that horizontal gene transfer may have shaped the functional capacity of the animal much more than previously suspected [14]. The analysis identified several thousand genes likely to be derived from nonmetazoan sources mostly from bacteria [15,16]. A second independent genome study of *H. dujardini* reported strong conflicts between the two assembled and annotated genomes although the biomaterial for both studies was taken from the same original stock culture [17]. Analysis of the second genome reference for *H. dujardini* indeed suggested a very low level of horizontal gene transfer. Lately, these studies were complemented by a similar analysis in a second tardigrade species, *Ramazzotius varieornatus* [18]. The authors leveraged their high-quality genome sequence of *R. varieornatus* and could show that only a small proportion of the gene coding complement represents putative foreign genes. The study further showed that the species (selectively) lost several members of pathways that promote stress damage (e.g., peroxisomal oxidative pathway, stress responsive pathway) during hypoxia, genotoxic or oxidative stress but simultaneously display expansion of gene families related to ameliorating damage (e.g., superoxide dismutases). A close examination of gene expression profiles during dehydration and rehydration by the authors revealed only minor differences between the two states. Additionally, the study identified a tardigrade-unique DNA-associated protein that, when transferred to human cell culture, suppresses DNA damage and promotes viability following irradiation. In summary, the study suggested that a) tardigrades can enter a dehydrated state without a massive transcriptional turnover and b) that the genome provides mechanisms that prevent, extenuate or protect against damage caused by extreme environmental conditions. Lately, a comparative study of the tardigrades *H. dujardini* and *R. varieornatus* revealed contrasting gene expression responses to anhydrobiosis [19]. While *H. dujardini* experienced a major transcriptional turnover, *R. varieornatu*s showed only limited regulation when switching to anhydrobiosis. The study further confirmed that the *H. dujardini* genome encodes only for a few horizontally transferred genes. Surprisingly, some of these seemed to be involved in the entry of anhydrobiosis. A whole-genome molecular phylogeny found more evidence for a Tardigrada+Nematoda relationship than the previously supported Tardigrada+Arthropoda relationship but also argued that a full genome sequence of representatives of Onychophora, more divergent Tardigrada and basally arising Nematoda would be required to fully address the tardigrade placement.

Here, we present an early draft genome sequence of the eutardigrade, *M. tardigradum* [20]. We have chosen this species because it is among the most stress resistant tardigrades [17]. Specimen even survived the exposure to space in low earth orbit [21]. Furthermore several transcriptomic [22–24], proteomic [25,26] and metabolomic [27] studies have already been performed within this species. Furthermore, this species is only related to the previously sequenced tardigrades. Its genome sequence not only enabled us to derive a more general view on the mechanisms of stress resistance. It also allowed us to corroborate the phylogenetic position of the tardigrades in the tree of life and, based on these results, to identify gene loss as a major evolutionary trend in both tardigrades and nematodes.

## Results

### A draft genome of *M. tardigradum*

The assembled genome of *M. tardigradum* comprises 75.1 Mb, in good agreement to the results of a flow cytometry based determination of 73.3 ±1.8 Mb (see Suppl. Fig. 1). Overall, 6654 contigs were assembled with a contig N50 size of 50 kb. The assembly was validated with two approaches. First, a prediction of 248 core eukaryotic genes with CEGMA [28] revealed a 96 % completeness of the genome. Second, a prediction of near-universal single-copy orthologs with BUSCO [29] was used to benchmark the *M. tardigradum* genome against three different lineages. Benchmarking against a nematode specific BUSCO set revealed a completeness of 41 % while benchmarks against an arthropod-specific and a metazoan-specific set revealed a completeness of 81 % and 82 % (see Fig. 1; Suppl. Table 1). Only 1.23 % of the assembly was classified as repetitive or low-complexity. Based on a metazoan repeat library, 1271 DNA transposons (mostly hobo-Activator and Tc1-IS630-Pogo) and 2033 LTR elements (mostly BEL/Pao and Gypsy/DIRS1) were identified. The integrative gene annotation approach predicted 19,401 protein coding genes. 1684 genes were putatively derived through tandem duplication while 43 genes probably originated through segmental duplication when compared to *H. dujardini* and *R. varieornatus*. The subsequent functional annotation found homologs for 12,518 genes (65 %) while 7534 had an ortholog within the reference species from Ensembl Metazoan Release 34 [26], *R. varieornatus* or *H. dujardini*. Based on the curated gene set, 10,966 genes were functionally assigned [30] to either a protein family, protein domain or a functional site, excluding low-complexity, transmembrane and coiled-coil assignments. 7357 genes had at least one associated gene ontology term. Out of 19,401 protein coding genes 261 were potentially derived through horizontal gene transfer while 665 were potential contaminations likely introduced during DNA extraction.

**Figure 1.**
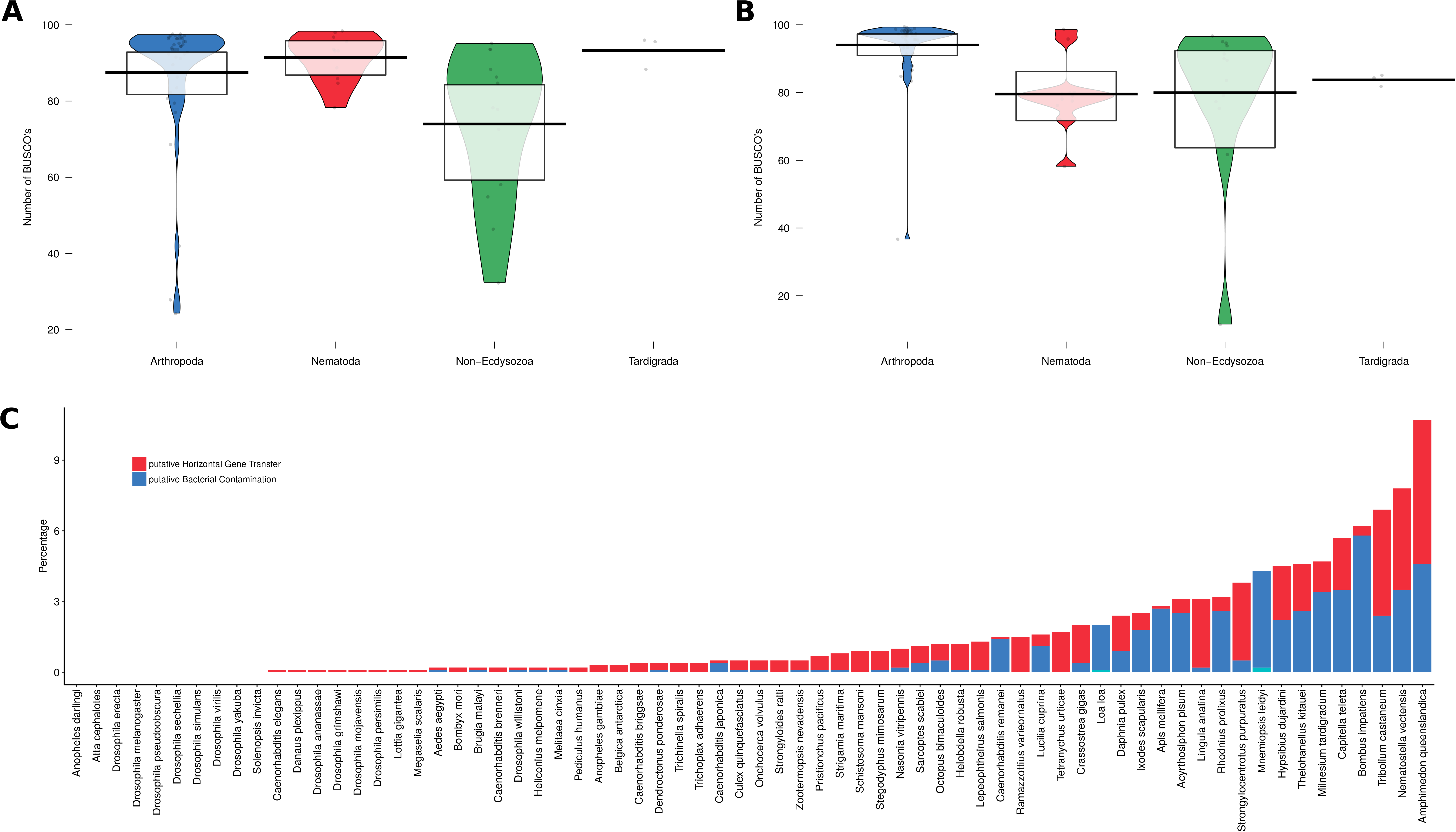
Genome completeness, contamination rate and putative percentage of horizontal gene transfer (HGT) across all Ensembl Metazoa Release 34 species and the three tardigrades. A) Genome completeness values of BUSCO and CEGMA. CEGMA completeness values show less spread within and between phyla. BUSCO completeness (using a set of metazoa BUSCOs) shows lower values for Nematoda, Non-Ecdysozoa and Tardigrada. B) Percentage of contamination-derived genes (e.g., bacterial proteins only on an individual genome sequence such as a contig) and HGT-derived genes (e.g., bacterial proteins on an individual genome sequence such as a contig surrounded by eukaryotic genes).

**Table 1.** Overview of components potentially involved in stress response identifiable in the three tardigrade genomes. First Block = Tardigrade-unique proteins present in the three tardigrade genomes. Second Block = Heat Shock Proteins present in the three tardigrade genomes. Third Block = Other stress-related proteins. *Abbreviations: MTAR = M. tardigradum; HDUJ = H. dujardini; RVAR = R. varieornatus.*

| Name | MTAR | HDUJ | RVAR |
| --- | --- | --- | --- |
| CAHS | 5 | 9 | 16 |
| SAHS | 0 | 5 | 13 |
| MAHS | 0 | 1 | 2 |
| Dsup | 0 | 0 | 1 |
| HSP70 | 12 | 69 | 13 |
| DNAJ | 42 | 43 | 33 |
| HSPB | 9 | 11 | 7 |
| HSPC | 3 | 2 | 2 |
| HSPD | 17 | 13 | 10 |
| HSPE | 1 | 1 | 1 |
| AOX | 1 | 1 | 1 |
| LEA | 1 | 1 | 0 |

### Phylogeny and hypothesis testing

Since the position of the phylum Tardigrada is still under discussion a phylogenomic analysis was performed to reconstruct the phylogeny underlying 56 Ensembl Metazoa species and the three tardigrades. Single copy clusters of orthologs with at least one tardigrade member were identified and used for phylogenomic reconstruction. Using all target sequences (selected species from Ensembl Metazoa Release 34 and the three tardigrades) 2245 of these clusters were detected. A Maximum Likelihood based phylogenetic reconstruction based on this supermatrix placed *M. tardigradum*, *R. varieornatus* and *H. dujardini* as the sister group of the nematodes (see Fig. 2). Next, we explicitly compared different hypotheses regarding the placement of the tardigrades, namely (i) as the sister group of the arthropods in a panarthropoda cluster, (ii) as the sister group of the nematodes grouping the tardigrades into the Cycloneuralia and (iii) as the outgroup to both arthropods and nematodes (see Fig. 4). Here, we used four different data sets. First, the super matrix was used to extract the per-site log-likelihood calculated by RaxML [31]. For each hypothesis, the approximately unbiased test as implemented in CONSEL [32] was performed. The same approach was carried out on (ii) the domain repertoire, (iii) the domain architectures and (iv) shared orthologous groups. As in the sequence based reconstructions, three tardigrades and 56 Ensembl Metazoa species were considered. In each case, the presence / absence of the feature was encoded in a binary matrix. For the super matrix and domain occurrences, the placement of the tardigrades as sister group to the nematodes had the highest rank (see Fig. 4). For shared orthologous groups, the placement as sister group to the arthropods had the highest rank while domain architectures ranked both placements equally. Taken together, the three different tardigrade genomes support, although not with full confidence, the tardigrades as members of a cycloneuralia cluster and reject their placement within the panarthropoda.

**Figure 2.**
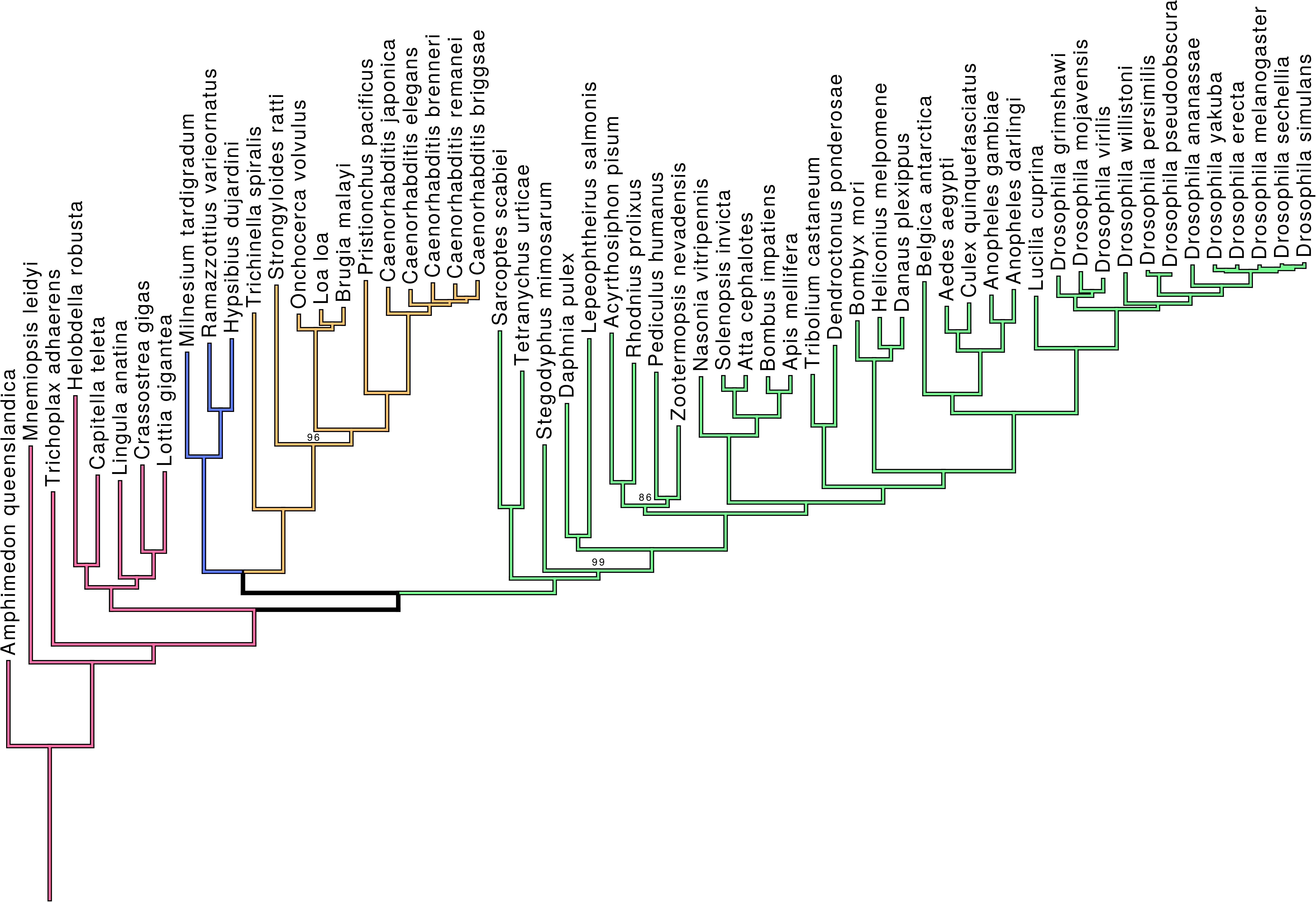
Maximum Likelihood based reconstruction of ecdysozoan phylogeny using a supermatrix approach. Only bootstrap support values with less < 100% support are shown. Additionally, all branches had a SH-aLRT support <= 80% [91]. Arthropods are placed as sister taxon to nematodes and tardigrades.

### Protein domain-ome comparison

The protein domain repertoire (domain-ome) of an organism establishes its functional capacity. Protein domains are evolutionarily conserved units usually with independent structural and functional properties [33–35]. They are widely distributed over all existing organisms [36] with some of them being universal and others being clade-specific. We tested whether the three tardigrade genomes shared patterns of domain family expansions, contractions or even total loss compared to the other 56 Ensembl metazoan species since these events could provide hints at which biological processes and molecular functions are most likely associated with tardigrade-specific traits. Protein domains were classified into Class I and II expansions as well as Class I and II contractions. Class I expansions and contractions are those where the query species experienced the highest or lowest protein domain occurrence count whereas Class II expansions and contractions indicated protein domains where the query species belongs to the group with the 5 % highest or lowest occurrence. Overall, 7939 protein domains were tested for contractions and expansions. In total, 169 protein domains tested significant for an expansion while 12 tested significant for a contraction (see Fig. 3, Suppl. Table 2) in at least one tardigrade. *M. tardigradum* showed 22 Class I expansions, 19 Class II expansions, 2 Class I contractions and 4 Class II contractions. *R. varieornatus* showed 36 Class I expansions, 26 Class II expansions, 0 Class I contractions and 3 Class II contractions while *H. dujardini* showed 35 Class I expansions, 31 Class II expansions, 2 Class I contractions and 1 Class II contractions. Only 12 domain expansions (ATPase_P-typ_cation-transptr_C, Hemopexin-like_repeat, Peptidase, metallopeptidase, Superoxide dismutase, copper/zinc binding domain, STAS_dom, BCS1_N, CLZ_dom, Glycine N-acyltransferase, N-terminal, LysM_dom, Nucleotide_cyclase, Peptidase M2, peptidyl-dipeptidase A, Peripla_BP_I; see Suppl. Table 2) overlapped between all three tardigrade species. Expanded domain sets from all three species were separately subjected to a gene ontology enrichment analysis (see Suppl. Table 3). Individual enrichments overlapped in 7 terms (cellular cation homeostasis, cellular ion homeostasis, multi-organism process, negative regulation of muscle contraction, response to organic substance, response to stimulus, response to stress). Surprisingly, all species showed a substantial amount of undetectable protein domains (*M. tardigradum*: 2239; *R. varieornatus*: 2146; *H. dujardini*: 2174) indicating potential loss events. The phylogenetic placement of tardigrades enabled us to examine the loss events in more detail. We used a Maximum Likelihood approach to reconstruct the domain repertoire of extinct ancestors. The approach enabled quantification of the domain loss using the phylogenetic tree while eliminating negative effects from contaminations or horizontal gene transfer events. We found that a major part of the losses happened before the split of tardigrades and nematodes followed by further tardigrade clade specific losses and only minor losses in the nematodes (see Fig. 5). No gene ontology terms were enriched in protein domains lost before the split of tardigrades and nematodes (Cycloneuralia losses) nor in protein (domains) specifically lost in tardigrades, arthropods or nematodes. Of note, the removal of putative contaminations and horizontally transferred genes had a larger impact on the numbers (e.g., nematode and arthropod specific losses). The assessment of domain family births revealed 2 domain families, namely SMC_ScpA (PF02616) and DUF612 (PF04747) gained before the split of tardigrades and nematodes. SMC_ScpA (PF02616) likely is an undetected contamination while DUF612 (PF04747) was only found in nematodes so far. Neither tardigrades nor nematodes showed further clade-specific gains.

**Figure 3.**
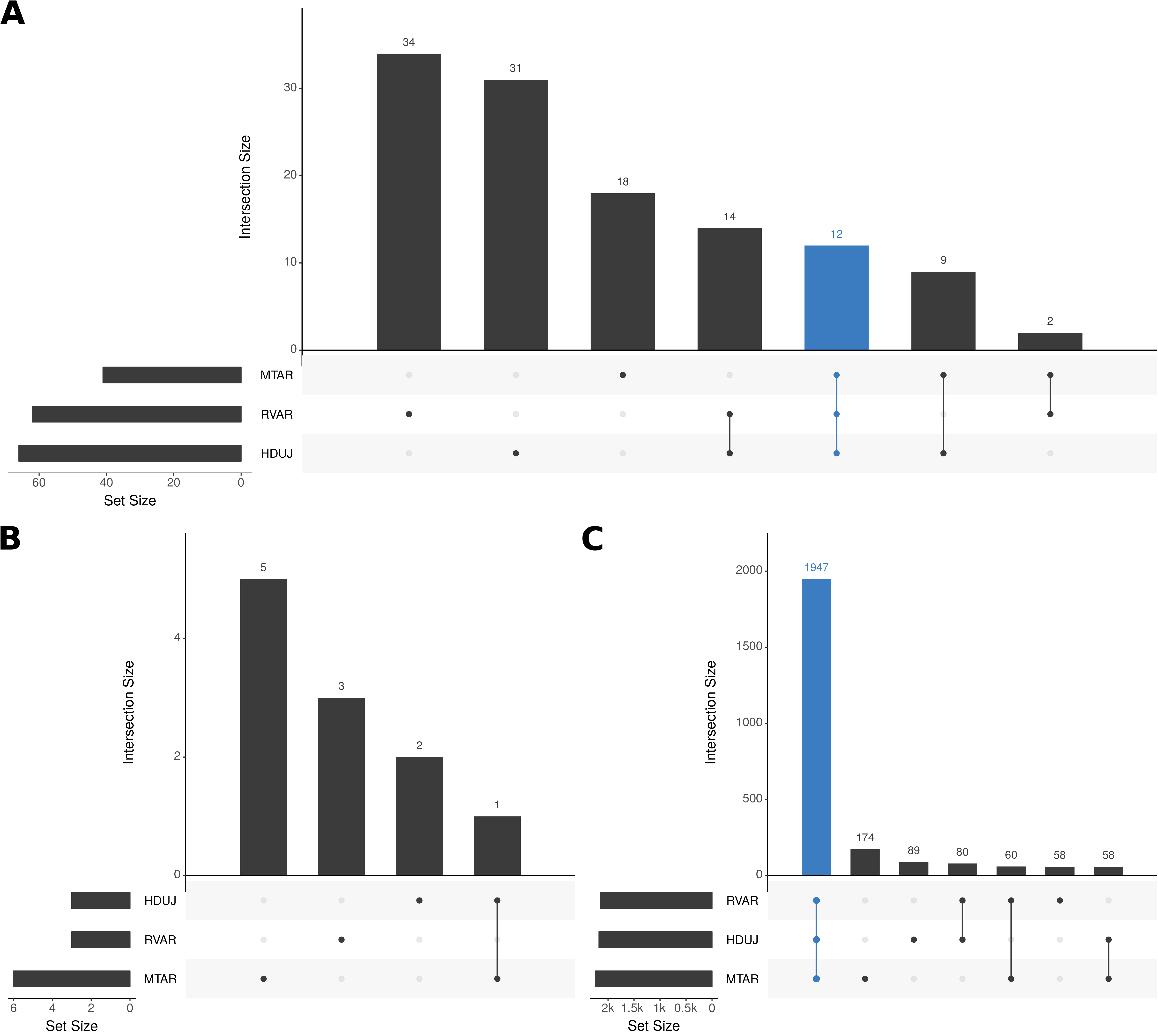
Hypothesis testing using the phylogenetic position of tardigrades. A) According to three different hypotheses, tardigrades are either the sister taxon to the arthropods (1), the nematodes (2) or the outgroup to both (3). B) Rank and probability of acceptance for each of these hypotheses for the four different datasets.

**Figure 4.**
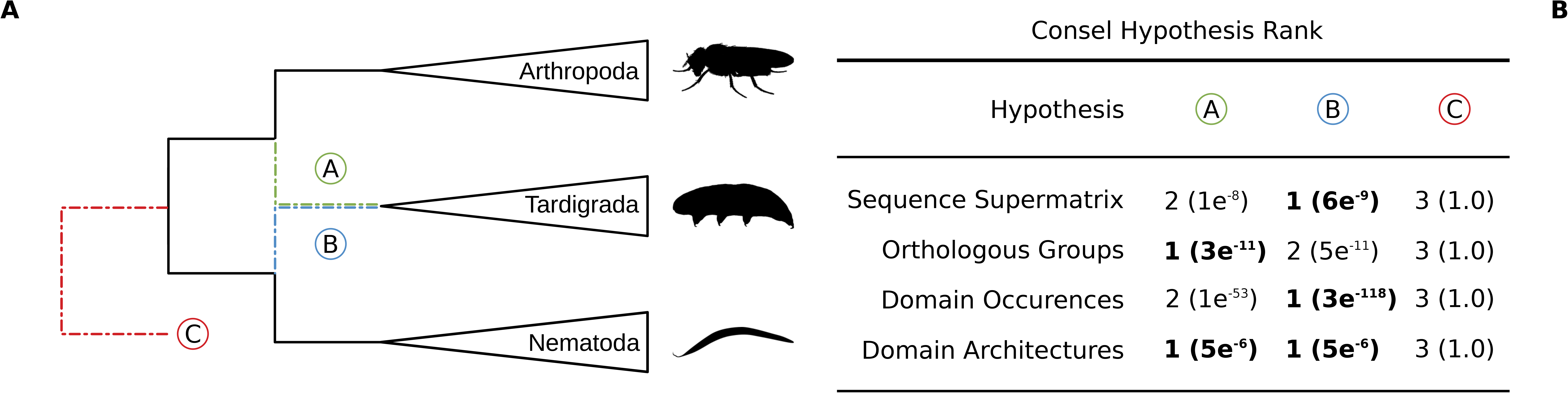
Intersection analysis of expanded, contracted and lost protein domain families in *M. tardigradum*, *R. varieornatus* and *H. dujardini*. Expansions were generally species-specific and only a small number was shared across all three species. Lost protein domains were largely shared across all three species. A) Expanded protein domains families, B) Contracted protein domain families and C) potentially lost protein domain families compared across the three tardigrade genomes. *Abbreviations: HD = H. dujardini, MT = M. tardigradum, RV = R. varieornatus*

**Figure 5.**
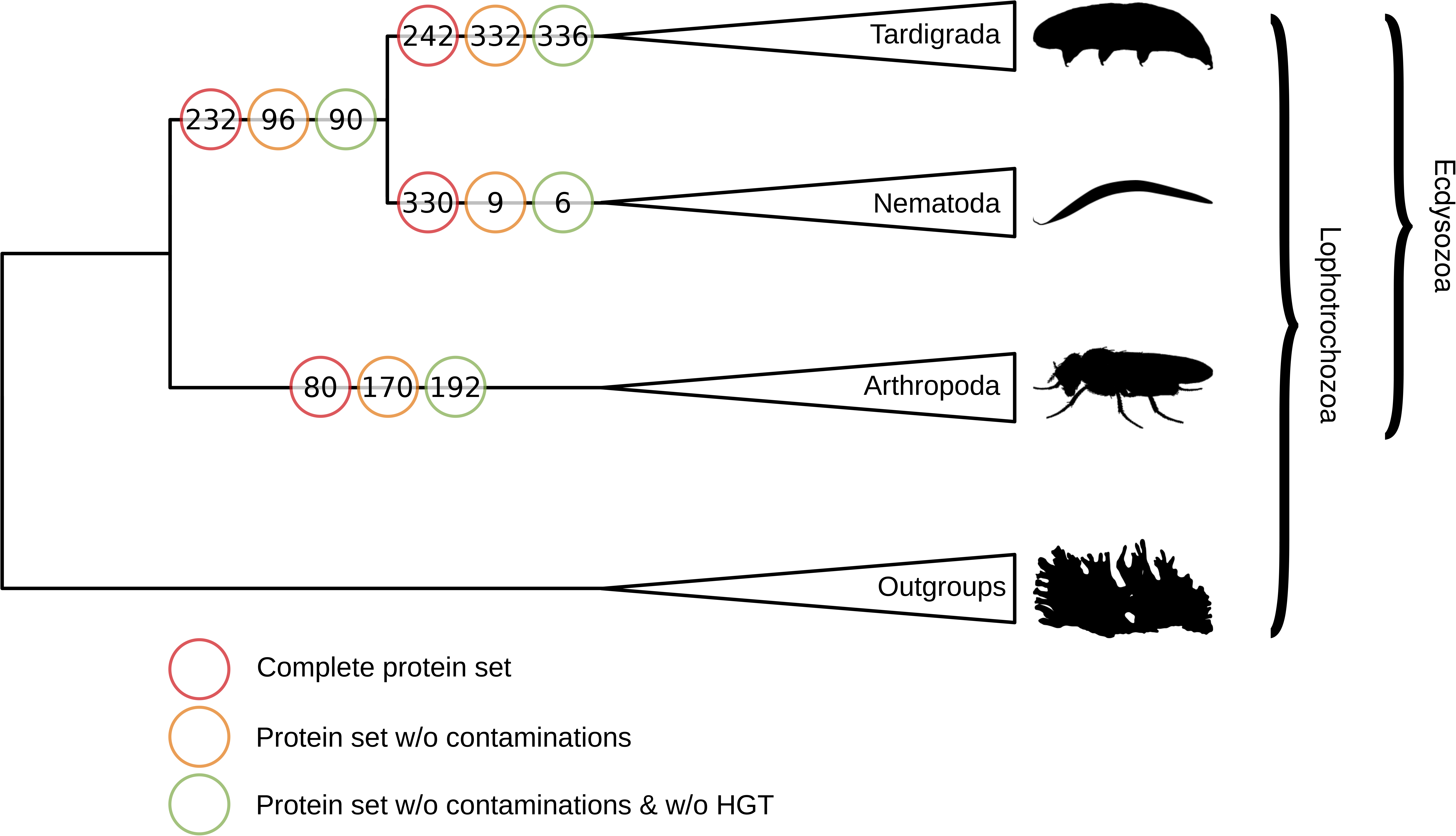
Domain loss in three Ecdysozoan lineages. The domain repertoire of ancestral species was reconstructed using Maximum Likelihood based on the accepted Cycloneuralia hypothesis (see Fig. 3, hypothesis 2). Colors: Red Circles = Loss estimates based on the complete protein sets; Orange Circles = Loss estimates based on protein sets without potential contaminations; Green Circles = Loss estimates based on the complete protein sets without potential contaminations and horizontally transferred genes

**Table 2.** Overview of components potentially involved in the trehalose metabolism identifiable in the three tardigrade genomes. First Block = Trehalose Synthesis. Second Block = Trehalose Degradation. *Abbreviations: MTAR = M. tardigradum; HDUJ = H. dujardini; RVAR = R. varieornatus.*

| Name | Description | MTAR | HDUJ | RVAR |
| --- | --- | --- | --- | --- |
| PF00128 | Alpha amylase, catalytic domain | 5 | 10 | 1 |
| PF00534 | Glycosyl transferases group 1 | 4 | 4 | 1 |
| PF00982 | Glycosyltransferase family 20 | 0 | 0 | 1 |
| PF02358 | Trehalose-phosphatase | 0 | 0 | 1 |
| PF03632 | Glycosyl hydrolase family 65 | 7 | 3 | 1 |
| PF01204 | Trehalase | 11 | 1 | 1 |

### Revisiting stress tolerance mechanisms

The enormous stress tolerance combined with the capability to undergo anhydrobiosis is arguably one of the most prominent features of tardigrades. Still, the underlying mechanisms are not fully clear. Even comparing the genomes of *H. dujardini* and *R. varieornatus* did not result in a consistent picture [19]. Rather, it was suggested, that already these closely related tardigrades utilize different strategies. The genome of *M. tardigradum* enabled us to revisit current hypotheses and to contrast the two Hypsibiidae genomes with the genome of a third more diverse eutardigrade from the family Milnesiidae. During the last years, mainly two strategies for the conservation of cellular structure in anhydrobiosis have been discussed. One of the proposed mechanisms is the accumulation of a specific sugar, trehalose. Indeed, the genomes of all three tardigrades encode at least one biosynthetic pathway from D-glucose to trehalose (see Table 2). The second hypothesis highlights the importance of late embryo abundance (LEA) proteins to stabilze cellular structures. Still, we identified the typical LEA protein domains in only two of the tardigrades (*M. tardigradum*: 1, *H. dujardini*: 1). Contradictory, 10 such proteins in *R. varieornatus* were reported earlier [18]. However, closer inspection of these proteins revealed that they did not contain typical LEA protein domains. Rather, some contained DUF883 (PF05957), a domain that frequently appears in proteins also containing LEA_4 (PF02987) protein domains. In addition to these most prominent hypotheses, a role of heat shock proteins (HSPs) in anhydrobiosis has been suggested. As these assist in protein folding and can refold denatured proteins, they could provide a self-evident mechanism to repair damage arising in anhydrobiosis. Indeed, all typical HSP protein families are encoded in the three tardigrade genomes (Table 1). Still, none of the families is significantly expanded in comparison to the other metazoan species. In addition to these hypotheses, Hashimoto et al. suggested the importance of several tardigrada-unique genes in the genome of *R. varieornatus* which are constitutively expressed and associated with stress tolerance. These included the previously identified Cytoplasmic Abundant Heat Soluble (CAHS) and Secretory Abundant Heat Soluble (SAHS) proteins as well as a newly characterized DNA Damage suppressor (Dsup). Based on the annotated *R. varieornatus* templates of these proteins we searched the genomes of *H. dujardini* and *M. tardigradum* using a profile and a homology based strategy. We found that all species encode several CAHS (see Table 1) but only *H. dujardini* and *R. varieornatus* encode SAHS proteins. Neither *M. tardigradum* nor *H. dujardin* has any homolog of Dsup.

The direct genome comparison of *H. dujardini* and *R. varieornatus* suggested that extensive loss in in the peroxisome pathway and in stress signaling pathways happened in both species and that loss of these resistance pathways may be associated with anhydrobiosis. We also searched for components that might act as reactive oxygen species (ROS) production suppressors. Surprisingly, we found an alternative oxidase (AOX) in each of the three tardigrades. AOX is erratically found in metazoan species but frequently present in many plants and bacteria. It can directly suppress ROS production and indirectly change the energy status of the cell through the alternative respiratory pathway [37]. In summary, the erratic distribution of tardigrade-unique proteins underlines their variable relevance for stress tolerance. Thus, there might indeed be a role of alternative pathways (e.g., ROS suppression through AOX) that are yet unexplored in tardigrades. Still, both facts support the notion that the phylum Tardigrada is a rich source of new protection genes, pathways and mechanisms involved in stress tolerance.

## Discussion

The remarkable stress tolerance of tardigrades has fascinated scientists and nonscientists alike. Accordingly, the publication of the first genome sequence of a tardigrade, *H. dujardini*, generated large interest. Still, the results presented in the original publication have been defeated largely [14–16]. Here, the genome of another tardigrade, *R. varieornatus* was of substantial importance. Both Eutardigrada species are closely related and belong to the family of Hybsibiidae. However, the phylum Tardigrada consists of more than 1000 species with a huge phenotypic and presumably also genetic variability. Thus, clade-wide insights from comparative genomics are limited to the species spectra analysed. Here, we present an early draft genome of a Milnesiidae species, *M. milnesium*. Although fragmented, it enabled us to test current hypotheses regarding the molecular mechanisms of tardigrade stress tolerance of tardigrades in a wider scope.

More than 40 years ago, water replacement and vitrification were suggested as core mechanisms for stabilization of cellular structures [37]. According to this hypothesis, water is replaced by other biomolecules, resulting in a glass-like state of the cell. Mainly two types of molecules enabling this transition, sugars [38,39] and late embryo abundant (LEA) proteins [40,41] have been described. An analysis of the trehalose biosynthesis and metabolism pathway showed that necessary components to generate and reconvert trehalose are indeed present in all three tardigrades. Although small amounts of trehalose were found *R. varieornatus* and *H. dujardini*, no evidence of the relevance for anhydrobiosis was found in *M. tardigradum* [10,42]. This could indicate independent adaptations in different branches of the tardigrades. LEA proteins were first detected in plants [43]. Later, these proteins were shown to be of importance for anhydrobiosis in nematodes [40] and arthropods [44]. With a phylogenetic position between these two phyla, tardigrades might also leverage these proteins to prevent protein aggregation. Still, genes encoding for typical LEA domain could be identified in only two of the tardigrades (*M. tardigradum* and *H. dujardini*).

As further candidates, heat shock proteins have been suggested. As they assist in protein folding and can refold denatured proteins, they could provide a self-evident mechanism to repair damage arising in anhydrobiosis. Still, the relevance of heat shock proteins for tardigrades is controversial. HSP70 expression is increased at rehydration [12] but not increased in desiccated animals [12,45]. Directly comparing different variants of HSPs revealed complex patterns [46,47]. We found several HSP-related protein domains significantly altered in each of the three tardigrade species. Thus, expansions previously published for the *R. varieornatus [17]* genome might be assigned to the whole clade.

The emergence of ROS is of considerable danger for a cell, as it can damage all cellular components. Already a challenge for a ‘standard’ cell, this problem increases dramatically when a cell desiccates. Accordingly, genes involved in the reduction of ROS are up regulated upon entry into anhydrobiosis [48]. A screen for protein domains related to ROS production and scavenging unexpectedly revealed that all three tardigrade genomes code for an alternative oxidase (AOX, PF01786). This protein can lower the internal production of ROS at the mitochondria [49,50]. While common in bacteria, plants and fungi, AOX was thus far only found in a few metazoan species, mostly living in salt water. This includes the bdelloid rotifer *Adineta vaga* which is also capable of anhydrobiosis [51] respectively cryptobiosis. In addition to the inactivation of ROS, avoiding its emergence would be a complementary strategy. The presence of AOX proteins might indicate an overlooked mechanism for anhydrobiotic metazoa and tardigrades as the so far first terrestrial animal utilizing this mechanism for ROS defense.

The genome of *R. varieornatus* was the first that revealed tardigrade-unique proteins involved in stress response. Hashimoto et al. (2016) discovered a protein that shields DNA from radiation (Dsup). Indeed, expression of this protein in human cells gave them a survival advantage. Yamaguchi et al. (2012) further discovered genes constitutively expressed and associated with stress tolerance like Cytoplasmic Abundant Heat Soluble (CAHS) and Secretory Abundant Heat Soluble (SAHS) proteins as well as Mitochondrial Abundant Heat Soluble (MAHS) proteins <u>(Yamaguchi et al. 2012)</u>. Surprisingly, genes coding for these proteins are absent in *H. dujardini* and *M. tardigradum*. In summary, the scattered distribution of MAHS-like, SAHS-like and tardigrade-unique proteins like Dsup, suggests species or at least class specific adaptations towards typical tardigrade traits.

The early draft genome presented here also provided new data to address the phylogenetic placement of the tardigrades, which is still discussed controversially. Mainly two hypotheses exist, placing tardigrades either as the outgroup of the arthropods building the Panarthropoda clade [53–58], or together with the nematodes as a member of the Cycloneuralia [59–64]. We improved the first phylogenetic analysis based on the genomes of *R. varieornatus* and *H. dujardini* first by including *M. tardigradum* to rely on a broader taxonomic range and second by including a wider range of species. The resulting tree reproduces the results of the previous analysis and robustly places the tardigrades as a sister group of the nematodes with the arthropods as outgroup. Still, phylogenetic reconstructions only report the most likely tree and will not provide a comparison to other tree topologies. To address this, we explicitly tested the current taxonomic hypotheses based on different datasets (sites in the supermatrix, orthologous groups, domain presence and domain architectures). This approach allowed us to provide further statistical support for each hypotheses. The hypothesis test on the supermatrix sites as well as the presence/absence of domain families supported the placement of the tardigrades as sister group to the nematodes consistent with the reconstruction of the tree. Testing shared domain architectures ranked none of two hypotheses (Tardigrada+Arthropoda and Tardigrada+Nematoda) highest while testing shared orthologous groups reversely ranked the Tardigrada+Arthropoda hypotheses highest. Still, we conclude that our analyses most strongly support the Cycloneuralia hypothesis.

With a reliable phylogenetic placement of the tardigrades at hand, we were able to reconstruct evolutionary events within the tree of the ecdysozoa. To this end, we used a maximum likelihood reconstruction of the ancestral nodes within the tree. We found, that the reduction of the domain repertoire starting from a complex Ur-ecdysozoan was a major trend in the evolution of the last common ancestor of nematodes and tardigrades was. This was followed by further lineage and species-specific losses, especially in tardigrades (see Fig. 5). Thus, the evolution of nematodes and tardigrades recalls the general trend of reduction already observed at the base of the Bilateria [65]. Notably, the reconstruction of evolutionary events was heavily influenced by contaminations and genes potentially derived through horizontal gene transfer. A more robust and contiguous genome reference for *M. tardigradum* and further support from transcriptomic, proteomic or other molecular experiments is necessary to ultimately link the adaptive genomic footprints to the unusual physiological capabilities of *M. tardigradum and* the tardigrades in general.

## Methods

### Animal culture

Tardigrade specimens of *M. tardigradum* Doyere 1840 [20] (Eutardigrada, Apochela), cultured in the laboratory for a decade, were used to study the genome. Originally, they were collected from dry moss in Tübingen, Germany. The carnivorous tardigrade species was reared in plastic culture dishes on a small layer of 3 % agar, covered with Volvic^TM^ water (Danone Waters Deutschland, Wiesbaden, Germany). Rotifers of the species *Philodina citrina* were provided as food twice a week. The cultures were maintained in an environmental chamber at 20 °C using an artificial light source with a 12 h light, 12 h dark cycle. For the DNA/RNA extraction exuvia with eggs and embryos were collected and cleaned by five washing steps with Volvic^TM^ water. Subsequently, they were placed separately in a 24-well plate until they hatched. Juveniles were transferred into a reaction tube, frozen in liquid nitrogen and stored at -80°C.

### Genome size estimation

The genome size of *M. tardigradum* was estimated using flow cell cytometry*. Drosophila melanogaster* was used as standard [67]. A culture of *M. tardigradum* was washed (4 times, M9 buffer) and placed into modified Galbraith’s buffer. Nuclei were released with a tissue grinder (Kontes Dounce tissue grinder, “A” pestle) and filtered to a 30 µm Nylon mesh. The same procedure was carried out with a single head from *D. melanogaster* female. The nuclear suspension was stained with propidium iodid (PI) for 2 h and measured immediately with a FACScalibur flow cytometer (Becton Dickinson, USA) and analyzed with CellQuest Pro version 6.0. PI-positive cells were gated and fluorescence intensity was analyzed in FL2-H channel and displayed on a linear scale (Supplementary Figure S1A). Non-stained cells served as a negative control (Supplementary Figure S1C). The whole procedure was benchmarked by comparing *D. melanogaster* (Supplementary Figure S1B) and *Apis mellifera* (data not shown) [68]. Results indicated an error of 5 %.

### Genome and transcriptome sequencing

DNA was extracted from approximately 1000 freshly hatched (to avoid bacterial contamination) animals using the Qiagen DNeasy kit according to the manufacturer’s instructions for animal tissues (spin column protocol). Animals were washed five times in RNAse/DNAse-free water, resuspended in Qiagen buffer ATL and disrupted with a FastPrep- 24 homogenizer (MP Biomedicals for 2 x 30 s at 4 m/s). Following overnight incubation in buffer ATL and proteinase K at 56 ºC, samples were treated with RNAse A and purified on a spin column. RNA was extracted from a similar sized animal culture as for the DNA. Animals were disrupted as described above, resuspended in buffer RLT, and RNA was extracted using the Qiagen RNeasy kit. Ribosomal RNA was depleted using the RiboMinus Kit for RNA-seq (Invitrogen) and reverse transcribed using random hexamers (Promega Im-Prom II Reverse Transcription System). cDNA was amplified using the GenomePlex Complete Whole Genome Amplification Kit (Sigma). The 95 ºC fragmentation step was omitted from the whole genome amplification, as RNA had been fragmented during homogenization. Bead libraries were prepared from DNA (1.8 µg) and cDNA (2 µg) using the GS FLX Titanium general library preparation kit (454 Life Sciences), followed by amplification using emulsion PCR with the LV emPCR kit (Lib-L) (454 life Sciences). Sequencing was performed on a 454 FLX instrument (454 Life Sciences). A second sequencing data set was produced from an additional batch of animals. DNA was extracted as above and subjected to whole genome amplification with Qiagen REPLI-g prior to sequencing. TruSeq DNA library prep and Illumina sequencing was carried out by GATC.

### Genome and transcriptome assembly

Genomic and transcriptomic reads were prepared by masking vector contamination and adapters using SMALT [69]. All read sets were compared against NCBI-nr using diamond [70]. The resulting alignments were prepared for MEGAN using daa-meganizer [71]. MEGAN was used to compute the lowest common ancestor for each read individually. Reads assigned only to the superkingdom Bacteria or Archaea were removed from the data set if there GC content was smaller than 25% or larger than 55%. Remaining genomic reads were assembled with Canu (release 1.3; errorRate=0.035, genome- Size=75000000, minReadLength=50, corMinCoverage=0, corMaxEvidenceErate=0.15, minOverlapLength=50, trimReadsCoverage=2) [72]. Transcriptomic reads were assembled using MIRA4 with accurate settings [73]. Genome completeness was validated with CEGMA [28] and BUSCO [29].

### Genome feature annotation

Known repetitive elements were annotated with RepeatMasker (v.4.0.4, species=metazoa) [74]. Coding genes were annotated with Braker1 (version 1.9; default parameters) [75] by combining de novo gene predictions and evidence alignments from ESTs. Evidence alignments were generated by aligning all ESTs against the genome using BLAT. Resulting alignments were converted into intron boundaries and passed to Braker1. The resulting proteins were functionally classified using homology and profile based methods. Protein families, domains and important sites were assigned using InterproScan5 (release 5.20; default parameters) [30] and the Interpro database (release 59.0) [76]. Signal peptides and transmembrane regions were predicted with SignalP (v.4.0) [77] and TMHMM (v2.0) [78]. Gene ontology terms and basic functional descriptions were assigned by lifting protein domain gene ontology annotations to their respective gene/protein. Segmental and tandem duplicates were detected using MCScanX [79].

### Protein Domain Expansions and Contractions

Significantly expanded and contracted protein domains (Pfam) were identified by comparing their occurrence in the three tardigrades (*H. dujardini* release 2.3.1 and *R. varieornatus* release Rv101 from http://ensembl.tardigrades.org) to all reasonable complete species present in the Ensembl Metazoa database (Release 34; see Suppl. Fig. 2) using a chi square test. The occurrence of a specific protein domain in the three tardigrades was compared to the occurrence of the same protein domain in each of the reference species individually. The number of all proteins associated with at least one protein domain was used as background for each species. The resulting p-values for each protein domain were combined into a weighted consensus p-value since they addressed the same null hypothesis, that a protein domain is not expanded or contracted significantly. For that, all p-values were z-transformed and a weighted consensus test was applied. The final weighted consensus p-value was adjusted using the Bonferroni method and considered significant at a level of 5%. Expansion and contractions were used to test for enriched gene ontology terms with dcGOR [80]. Enrichments were statistically verified with the hypergeometric test (Parent-Child algorithm). P-values were adjusted using Bonferroni’s method. All proteins domains found in the 56 species were used as background.

### Phylogenetic reconstruction and hypothesis testing

The phylogenetic reconstruction was carried out using 56 selected species present in release 34 of the Ensembl Metazoa database [81]. Putative contaminations and horizontally transferred genes in all species were identified by comparing their predicted proteins against the Ensembl Metazoa database (excluding the query species) as well as all bacterial RefSeq non-redundant proteins using diamond [70]. The resulting alignments were prepared for MEGAN using daa-meganizer [71]. MEGAN was used to compute the lowest common ancestor for each read individually. Proteins with bacteria as lowest common ancestor which were encoded on contigs only containing bacterial proteins were flagged as putative contamination. Proteins with bacteria as lowest common ancestor but flanked by at least two eukaryotic proteins were flagged as horizontally transferred. Proteins flagged as contamination nor as horizontally transferred were excluded from all analysis if not stated otherwise. Potential in-paralogs, orthologs and co-ortholog pairs were identified using orthAgogue [82]. The species phylogeny was reconstructed using IQ-TREE (version 1.5.5; - alrt 1000 -bb 1000 -bspec GENESITE -m TEST) [83] based on the ortholog groups generated with MCL [84]. Only single copy ortholog groups which contained at least one tardigrade and a minimum number of 12 species were considered. The maximum-likelihood tree was inferred using the edge-linked partition model in IQ-TREE [85]. Branch support was obtained with the ultrafast bootstrap method [86]. Substitution models were selected using ModelFinder [87]. Alternative phylogenetic hypothesis for the placement of the tardigrades were tested with RAxML [31] and CONSEL [32]. Testing was done using binary representations of the absence-presence matrices for protein domain families, domain architectures and orthologous groups. Domain architectures were defined on Pfam domain families [88]. Repetitive stretches of domains were collapsed and the order not considered. RAxML (-f A -m BINGAMMA -T 2) was used to calculated per-site log likelihoods for each of the alternative hypothesis. Test statistics for alternative hypothesis were calculated using the ‘approximately unbiased test’ implemented in CONSEL with 100 replicates.

### Protein domain loss estimation

Protein domain losses and gains were detected using the Tardigrada+Nematoda (Cycloneuralia) tree topology and the corresponding absence-presence matrices. Ancestral nodes were reconstructed using RAxML (-f A -m BINGAMMA -T 2).

### Identification of domains associated with stress resistance

SAHS/CAHS containing proteins previously identified in *R. varieornatus* were detected using a profile based approach. Template from *R. varieornatus* were aligned, the alignment manually curated and used to build a hidden Markov model (HMM) [89]. A reverse search of the model against *R. varieornatus* proteins was conducted and the results used to define an optimal inclusion e-value (CAHS = 1.2×10^−22^; SAHS = 5.1×10^∼40^). The final model was used to screen proteins from all species. Dsup homologs were identified by a simple protein blast (BLASTP) [90] against the complete set of 56 Ensembl Metazoa Release 34 species and the three tardigrades. HSPs were identified by using predefined Pfam protein domains (PF00011, PF00012, PF00118, PF00166, PF00183, PF00226). Trehalose biosynthesis and metabolism components were identified by using predefined Pfam protein domains (PF00128, PF00358, PF00534, PF00982, PF01204, PF02056, PF02358, PF02922, PF03632, PF03633, PF03636, PF09071, PF11941, PF11975, PF16657). Late Embryogenesis Abundant proteins (LEA proteins) were identified using predefined Pfam protein domains (PF00477, PF02987, PF03168, PF03242, PF03760, PF10714). AOX proteins were identified using the predefined Pfam protein domain (PF01786).

## Acknowledgements

We would like to thank Steffen Hengherr for the maintenance of the tardigrade culture and Thomas Hegna for the drawing of *D. melanogaster* used in Figures 3 and 5.

## Abbreviations

ROS: Reactive Oxygen Species
HSP: Heat Shock Protein
LEA: Late Embryo abundant
AOX: Alternative Oxidase
HGT: Horizontal Gene Transfer
HMM: hidden Markov model
CAHS: Cytoplasmic Abundant Heat Soluble
SAHS: Secretory Abundant Heat Soluble
MAHS: Mitochondrial Abundant Heat Soluble

## Accession Numbers

Raw sequencing data, the genome assembly and its gene annotation are deposited in EBI ENA under accession number PRJEB22082.

**Supplemental Figure 1.**
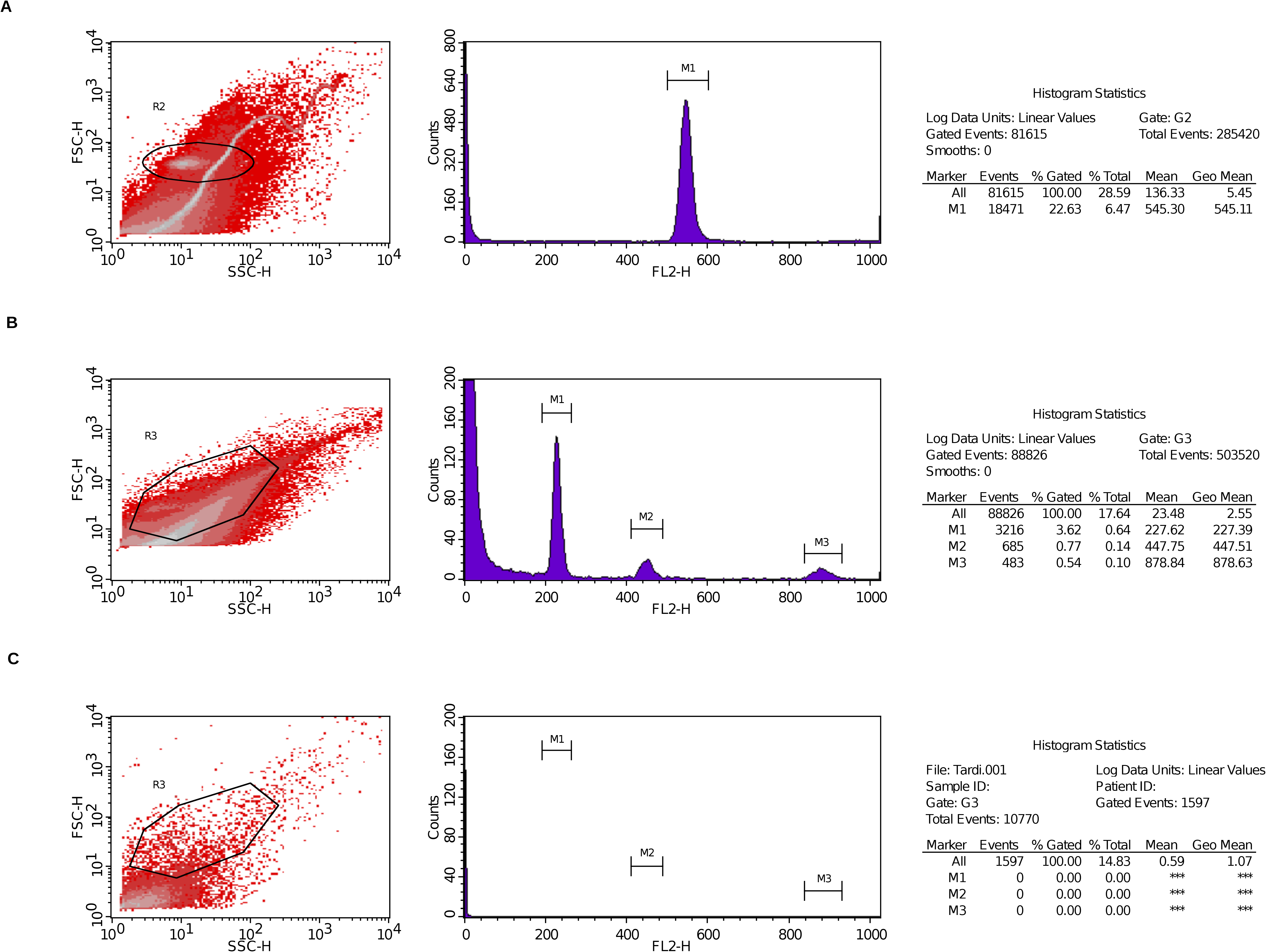
Genome size estimation for *M. tardigradum*. The histograms of relative DNA content were obtained after flow cytometric analysis of propidium iodide-stained nuclei. A) Stained nuclei from whole *M. tardigradum* animals; B) Stained nuclei from a single *D. melanogaster* head; C) Unstained nuclei from whole *M. tardigradum* animals. Marker M1 corresponds to the diploid genomes size in all samples. The ratio of M1 peak means (*M. tardigradum*: *D. melanogaster*) was equal to 0,41 and hence the 2C DNA amount of *M. tardigradum* was estimated to about 0,75 pg corresponding to a genome size of 73.3±1.8 Mb (SD was calculated from a cross comparison of *D. melanogaster* and *A. mellifera*, data not shown).

**Supplemental Figure 2.**
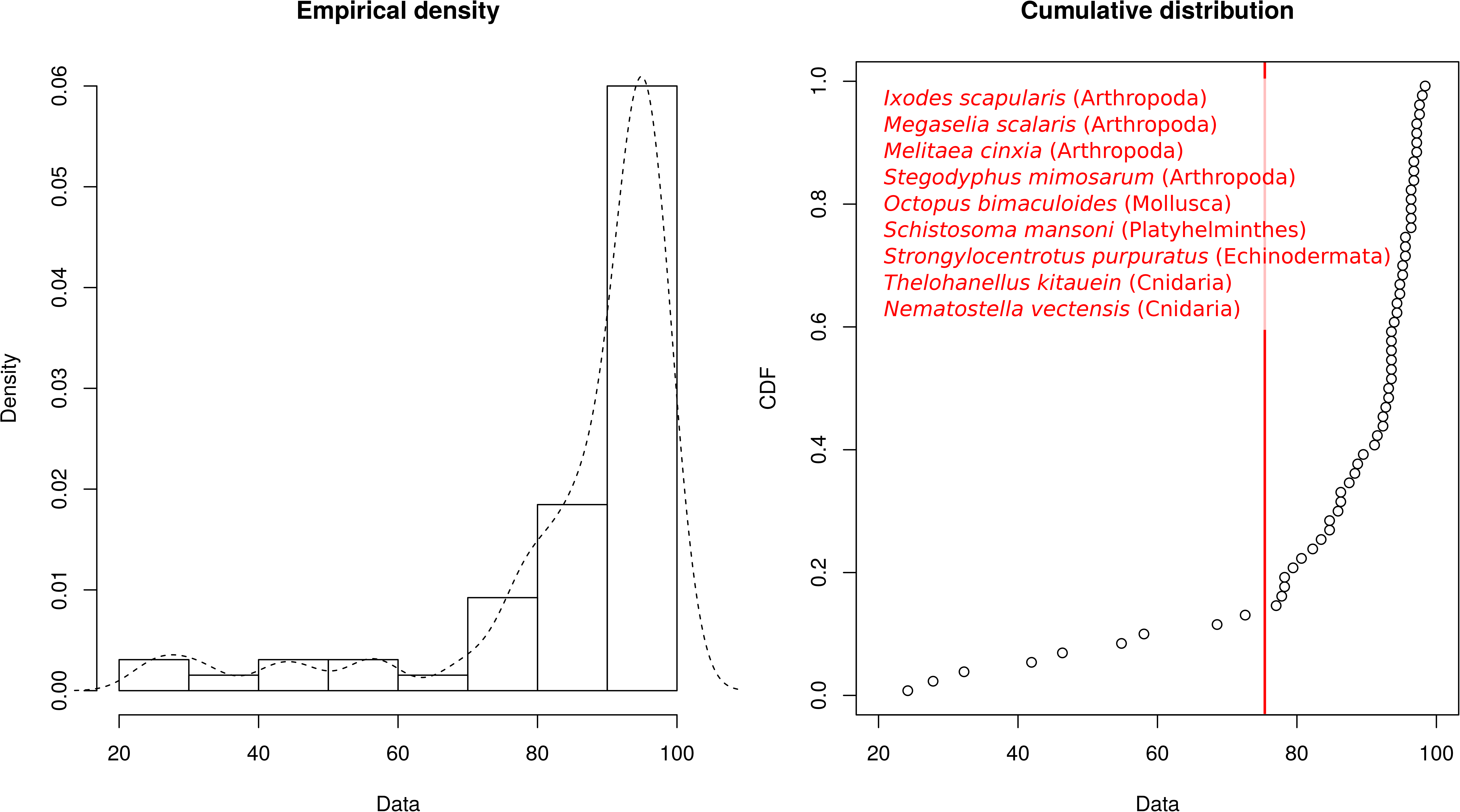
Ensembl Metazoa genome selection based on visual inspection of their completeness values. Empirical density and the cumulative distribution of CEGMA completeness values. 75% was chosen as the final threshold. species indicated in red were excluded based on the threshold.

